# Medulloblastoma Spatial Transcriptomics Reveals Tumor Microenvironment Heterogeneity with High-Density Progenitor Cell Regions Correlating with High-Risk Disease

**DOI:** 10.1101/2024.06.25.600684

**Authors:** Franklin Chien, Marina E. Michaud, Mojtaba Bakhtiari, Chanel Schroff, Matija Snuderl, Jose E. Velazquez Vega, Tobey J. MacDonald, Manoj K. Bhasin

**Author notes:** Co-First authors (Franklin Chien, Marina E. Michaud). Senior Authors (Tobey J. MacDonald, Manoj K. Bhasin). **Senior and Corresponding authors** Manoj K. Bhasin, MS, PhD, Aflac Cancer and Blood Disorders Center Children Healthcare of Atlanta, Health Sciences Research Building II, Room N320 1760 Haygood Drive, 3^rd^ Floor, Emory School of Medicine Atlanta, GA 30322; Tobey J. MacDonald, MD, Aflac Cancer and Blood Disorders Center Children’s Healthcare of Atlanta, Health Science Research Building I, Room E384 1760 Haygood Drive, 3^rd^ Floor, Emory School of Medicine Atlanta, GA 30322.

## Abstract

The tumor microenvironment (TME) of medulloblastoma (MB) influences progression and therapy response, presenting a promising target for therapeutic advances. Prior single-cell analyses have characterized the cellular components of the TME but lack spatial context. To address this, we performed spatial transcriptomic sequencing on sixteen pediatric MB samples obtained at diagnosis, including two matched diagnosis-relapse pairs. Our analyses revealed inter- and intra-tumoral heterogeneity within the TME, comprised of tumor-associated astrocytes (TAAs), macrophages (TAMs), stromal components, and distinct subpopulations of MB cells at different stages of neuronal differentiation and cell cycle progression. We identified dense regions of quiescent progenitor-like MB cells enriched in patients with high-risk (HR) features and an increase in TAAs, TAMs, and dysregulated vascular endothelium following relapse. Our study presents novel insights into the spatial architecture and cellular landscape of the medulloblastoma TME, highlighting spatial patterns linked to HR features and relapse, which may serve as potential therapeutic targets.

## Introduction

Medulloblastoma (MB), the most prevalent pediatric central nervous system (CNS) malignancy, is a biologically and clinically heterogeneous group of grade IV embryonal tumors of the posterior fossa, presenting significant clinical challenges. Advancements in genomic profiling have identified four primary molecular groups of MB: wingless (WNT) activated, sonic hedgehog (SHH) activated, Group 3, and Group 4, with the WNT subgroup associated with the most favorable outcomes and Group 3 the least favorable^1–3^. This categorization became the predominant subgrouping reflected in the most recent World Health Organization (WHO) 2021 Classification of CNS tumors^4^, along with a separate category for traditional histological classification.

Additional features such as Chang’s metastasis staging system, histological classification, and molecular alterations, such as *TP53* mutation, *MYC*- and *MYCN*-amplification, and isochromosome 17q further refine prognostication and segregate tumors to clinically high-risk (HR) and standard-risk (SR) disease with respect to response to therapy. Therapy includes maximally safe resection, craniospinal irradiation, and DNA-alkylating chemotherapies^5,6^. Despite multimodal therapy, 5-year overall survival remains 70-85% in those with SR disease and is lower for patients with HR defined by subtotal resection (STR), metastatic disease at diagnosis, or by molecular features as defined by the most current Children’s Oncology Group risk stratification classifications^7^. Approximately 20% of cases report STR, and 30% present with metastatic disease at diagnosis, resulting in lower 5-year progression-free survival^8^. The need for new therapeutic strategies targeting HR disease is underscored by the increased hazard ratios of 1.67 for STR and 1.45 for metastatic disease relative to non-metastatic and gross-total resection groups^8^.

The MB tumor microenvironment (MB-TME) plays a central role in MB disease progression and relapse. Thus, therapeutic strategies to target the components of the TME have emerged as the focus of multiple studies^9^. In recent years, investigations employing single-cell RNA-sequencing (scRNA-seq) have provided unprecedented insight into genomic and cellular heterogeneity of the MB-TME, uncovering subtypes of malignant, immune, and stromal cells^10,11^. As an immunologically cold tumor, the MB-TME harbors low proportions of tumor-infiltrating lymphocytes (TILs), along with brain-resident populations, supporting stromal components, and blood and lymphatic vasculature^9^. Tumor-associated microglia and macrophages (TAMs) are the most prevalent immune population and most prominent in SHH-activated disease^9^. TAMs can exhibit both anti- and pro-tumoral functions in the MB-TME as a result of polarization, wherein the M2 phenotype has been associated with tumor growth and progression^12^. On the other hand, decreased TAM abundance has been associated with poor survival in human and murine orthotopic models,^13^ with high percentages of M1 macrophages linked to favorable outcomes^14^. Tumor-associated astrocytes (TAAs) are a prominent cellular component in high-grade CNS tumors, implicated in tumor progression and metastasis^15^. In medulloblastoma specifically, TAAs have been recently found to increase tumor cell stemness, survival, and proliferation through the secretion of intercellular signaling molecules, including SHH, chemokine C-C ligand 2 (CCL2), lipocalin-2 (LNC2), and tumor necrosis factor alpha (TNF-α)^16,17^.

Malignant cells within the MB-TME display heterogeneous phenotypes, primarily defined by varying stages of neuronal differentiation and cell cycle progression. Notably, scRNA-seq studies by Hovestat *et al*. and Riemondy *et al.* have identified three primary transcriptional programs in MB cell states: undifferentiated progenitor, cell cycling, and neuronally differentiated, elegantly characterizing the inter- and intra-tumoral heterogeneity of MB. Distinct malignant cell phenotypes are known to drive disease progression, therapeutic responses, and patient outcomes. Platinum-based chemotherapies rely on the proliferative behavior of cancer cells, with therapy resistance driven by quiescent malignant subpopulations arrested in a reversible non-proliferative state (G0) termed dormant cancer cells^18,19^. These findings extend to CNS tumors, wherein quiescent states associated with disease progression have been identified in prominin-1 positive (PROM1^+^) glioblastoma cells, as well as in SRY-Box transcription factor 9 positive (SOX9^+^) cells in MYC-driven MB^20^.

While single-cell analysis has enabled new insights into the cellular composition of the MB-TME, a critical limitation to scRNA-seq is that the spatial organization of cell types in relation to one another and within tissue architecture is lost. While probe-based microscopy techniques can address this limitation, providing spatially resolved gene and protein expression data, they remain limited by low-throughput and cell-type localization extrapolated from limited markers. Spatial high-throughput methods are needed to identify TME cell composition, including dormant or sleeping cancer cells and their spatial positions. To address this gap, we present the first unbiased spatial sequencing of MB samples from patients across all molecular subgroups with diverse clinical characteristics. Our work provides spatial profiling of the diverse cellular architecture of MB and its associations with clinical features, identifying potential targets for therapeutic intervention.

## Results

### Characteristics of the clinical cohort for spatial profiling

Formalin-fixed paraffin-embedded (FFPE) blocks were obtained for sixteen tumor biopsy samples from fourteen individual patients with medulloblastoma and profiled by spatial transcriptomic sequencing using the Visium platform (**Figure 1a**). For all 14 patients, samples were obtained at initial diagnostic resection, with two additional patient-matched samples obtained at relapse after chemotherapy and radiation. The cohort includes all four major molecular subgroups: SHH-activated subtype (*n* = 6), WNT-activated subtype (*n* = 1), Group 3 (*n* = 2), and Group 4 (*n* = 5).

**Figure 1.**
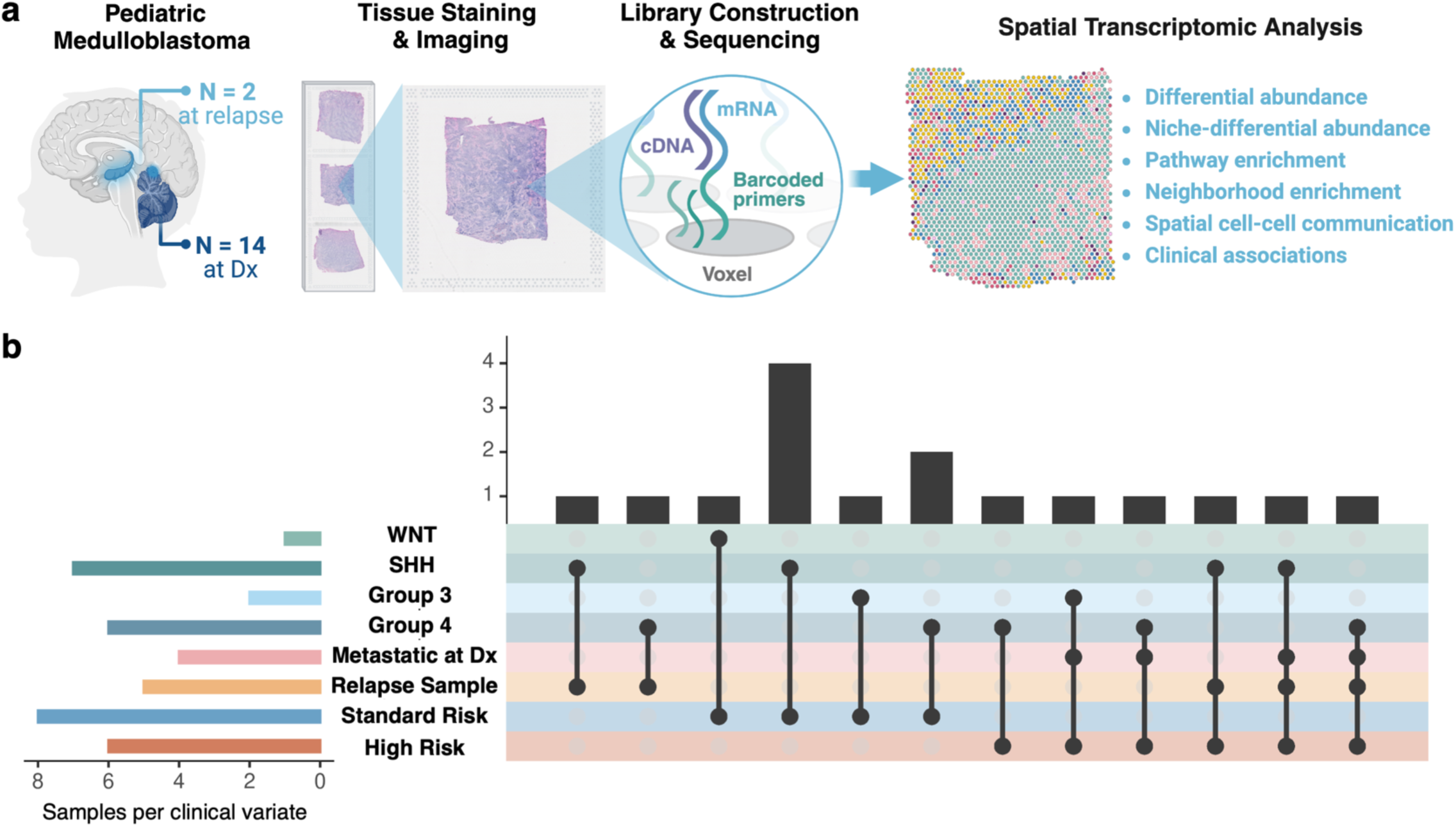
Overview of study design for mapping the spatial molecular landscape of Medulloblastoma. **(a)** Using the 10X Genomics Visium platform, spatial sequencing was performed on 16 formalin-fixed paraffin-embedded (FFPE) samples taken at diagnosis or relapse from pediatric patients with medulloblastoma. **(b)** The clinical cohort consisted of 14 patients representing a range of clinical characteristics, including metastatic disease, relapsed disease, and risk stratification. The cohort includes all molecular subgroups: SHH-activated subtype (*n* = 6), WNT-activated subtype (*n* = 1), Group 3 (*n* = 2), and Group 4 (*n* = 5). The cohort contains standard (n=8) and high (n=6) risk patients obtained at the time of disease diagnosis. The cohort contains paired samples from two patients collected at the time of disease diagnosis and relapse.

Among these patients, a range of demographics and clinical characteristics are present (**Figure 1b**, **Supplemental Material S1**). Of the cohort patients, 13 were male and 1 female, with racial backgrounds including non-Hispanic Black (*n* = 7), Hispanic White (*n* = 3), and Non-Hispanic White (*n* = 4) and ages ranging from 1 to 22 years old. Histologically, the samples included classical (*n* = 6), large cell/anaplastic (LC/A, *n* = 5), and desmoplastic/nodular (DN, *n* = 4) types. Four patients presented with metastatic disease (M+) at diagnosis, and the remaining had no evidence of clinical metastasis (M0). Three patients, representing SHH, Group 3, and Group 4 subtypes, displayed *MYC*- or *MYCN*-amplification. Of the two Group 3 tumors, one exhibited *MYC* amplification and M+ disease, while the second did not. Patients were treated with either a standard-(*n* = 5) or high-risk (*n* = 3) Children’s Oncology Group (COG) protocol or with a radiation-sparing regimen due to younger age-at-diagnosis (*n* = 6). Of those receiving radiation-sparing due to young age at diagnosis, three were classified as high-risk based on the presence of metastasis at diagnosis. The remaining had no high-risk features and were classified as standard-risk. No patients were high-risk based on subtotal resection.

**Table 1.** Characteristics of study tumor samples (n=16)

|  | Standard-Risk<br>at Diagnosis | High-Risk<br>at Diagnosis | At Relapse |
| --- | --- | --- | --- |
| **n (%)** | 8 (50) | 6 (37.5) | 2 (12.5) |
| *Age (mean (SD))* | 7.6 (6.6) | 9 (5.5) | 19.5 (0.7) |
| **Sex (%)** |  |  |  |
| *Female* | 1 (12.5) | 0 | 0 |
| *Male* | 7 (87.5) | 6 (100) | 2 (100) |
| **Race/Ethnicity (%)** |  |  |  |
| *Non-Hispanic Black* | 4 (50) | 3 (50) | 2 (100) |
| *Non-Hispanic White* | 2 (25) | 2 (33.3) | 0 |
| *Hispanic* | 2 (25) | 1 (16.7) | 0 |
| **Stage at Diagnosis (%)** |  |  |  |
| *M0* | 8 (100) | 2 (33.3) | n/a |
| *M+* | 0 | 4 (66.6) | n/a |
| **Molecular Classification (%)** |  |  |  |
| *SHH-activated* | 4 (50) | 2 (33.3) | 1 (50) |
| *WNT-activated* | 1 (12.5) | 0 | 0 |
| *Group 3* | 1 (12.5) | 1 (16.7) | 0 |
| *Group 4* | 2 (25) | 3 (50) | 1 (50) |
| **Histological Classification (%)** |  |  |  |
| *Classic* | 4 (50) | 2 (33.3) | 1 (50) |
| *Desmoplastic/Nodular* | 3 (37.5) | 1 (16.7) | 0 |
| *Large Cell/Anaplastic* | 1 (12.5) | 3 (50) | 1 (50) |
| **Molecular Features (%)** |  |  |  |
| *MYC/MYCN-amplification* | 0 | 3 (50) | 1 (50) |
| *TP53 Mutation* | 0 | 0 | 0 |
| **Outcomes (%)** |  |  |  |
| *Survival* | 6 (75) | 3 (50) | 0 |

### Spatial profiling characterizes malignant, glial, and stromal components of MB-TME

FFPE slides derived from patient biopsies were reviewed by a neuropathologist to ensure viable and representative areas of tumor tissue were selected for spatial profiling. The selected tissue sections were transferred to the Visium slide (10X Genomics Inc.) capture area, followed by H&E staining, permeabilization, ligation, cDNA library construction, and sequencing (**Figure 1a**). The spatial profiling captured 1,000–3,000 voxels per sample, with 2,524 median transcripts per voxel representing 16,238 unique genes in the dataset (**Figure S1**). Following normalization and integration, principal component and clustering analyses illustrate shared transcriptional profiles within the TME across molecular subgroups, with clusters representing voxels expressing predominant malignant or non-malignant cell type signatures (**Figure 2a**).

**Figure 2.**
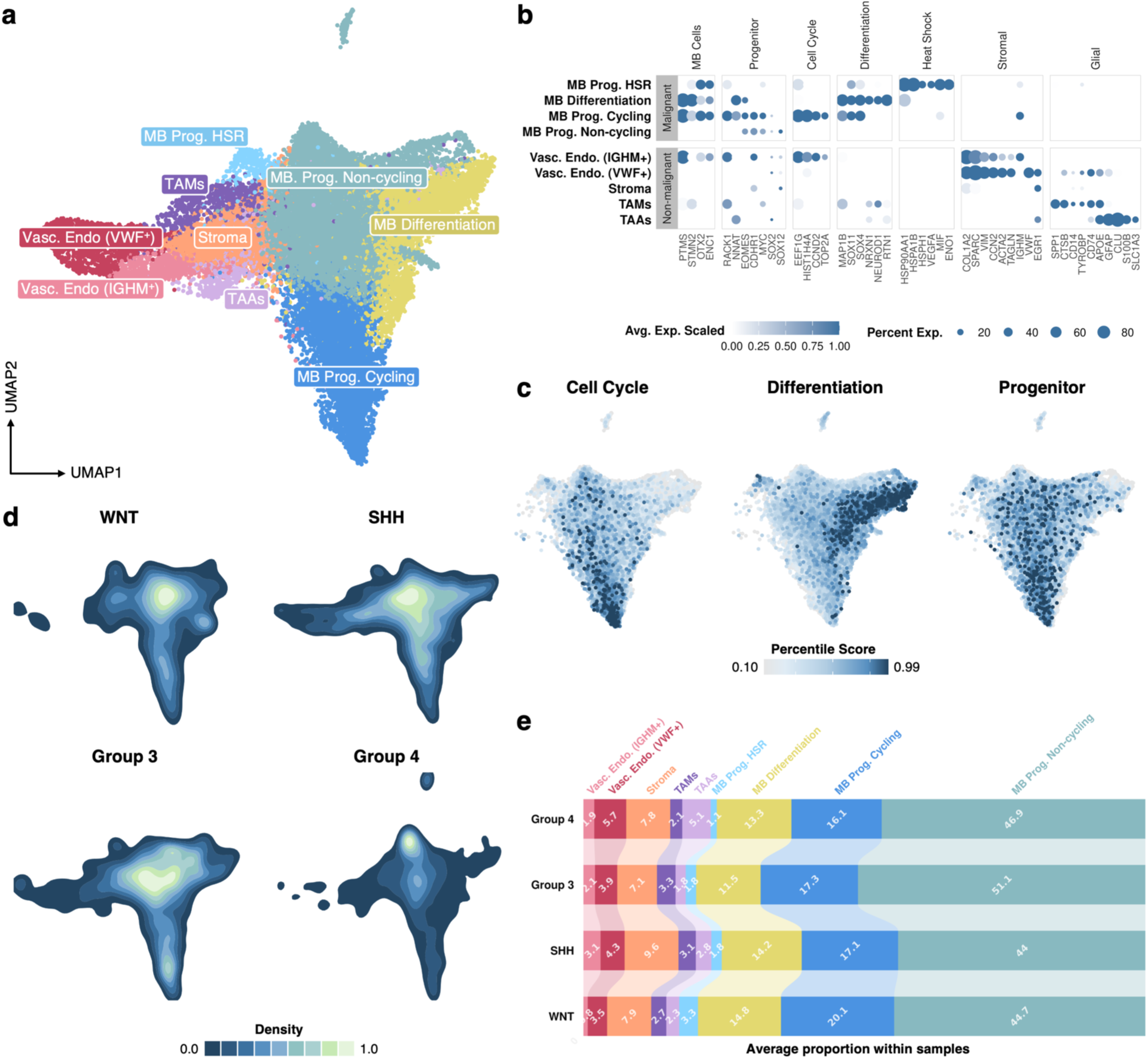
Spatial profiling characterizes malignant, glial, and stromal components of MB-TME. **(a)** UMAP visualization of K-NN clustering of voxels based on their transcriptomes yielded nine clusters, including four clusters with predominant medulloblastoma tumor markers and five clusters exhibiting predominant stromal signatures. The clusters are colored based on major cell types. **(b)** Dot plot displaying the expression markers related to major cell types and states present among clusters. The blue color intensity represents gene expression, and the size of the dot represents the percentage of voxels expressing each gene in different cell type clusters. **(c)** Feature plots for medulloblastoma clusters illustrating the enrichment of transcriptional programs relating to neuronal progenitors, cell cycle progression, and neuronal differentiation, aligning with previous medulloblastoma single-cell studies reported by Hovestadt *et al*. and Riemondy *et al*. The signatures for the transcriptional programs are shown in Table S1. **(d)** Density plots illustrating the differential enrichment of the cell types and states between MB molecular subgroups. **(e)** Bar plot displaying the average composition of clusters between molecular subgroups.

Among non-malignant cell types, clusters enriched for makers of TAAs, TAMs, stroma, and vascular endothelium were present (**Figure 2b**). Interestingly, two clusters of vascular endothelium emerged: one enriched for markers associated with immune infiltration (*IGHM*, *IGHD*) and the other for markers of endothelial activation and potential dysregulation (*VWF*, *EGR1*). Within the TAM-associated cluster, markers of both monocyte-derived macrophages and resident microglia were present, though further subclustering did not resolve these populations. Notably, these TAMs highly expressed the cytokine osteopontin (*SPP1*). SPP1^+^ TAMs have been implicated in tumor progression across several cancers, including brain cancers, but their role in medulloblastoma remains unexplored ^21,22^. In our dataset, *SPP1* expression was significantly higher in non-metastatic compared to metastatic samples; however, increased *SPP1* expression was also observed following relapse in matched samples (**Figure S2**).

Clusters exhibiting malignant markers revealed distinct MB subpopulations corresponding to those previously reported in single-cell transcriptomic analyses^10,11^ (**Figure 2c**). Specifically, previously described transcriptional programs representing MB progenitors and differentiating MB cell phenotypes were enriched in these clusters (see Methods), with progenitor cells further subdividing into cycling and non-cycling based on cell cycle gene enrichment. Additionally, we identified a unique cluster of malignant cells with elevated expression of heat-shock response (HSR) proteins. HSR proteins, including the HSP90 and HSP70 family protein genes observed, have a well-documented role in tumor progression, including brain cancers, through cell survival regulation^23^. Their expression in MB has been previously reported as potentially promising therapeutic targets warranting further investigations^24,25^. Notably, Group 3 disease, associated with the poorest clinical outcomes, displayed increased density within the cycling MB progenitor cluster (**Figure 2d**). In contrast, Group 4 disease exhibited increased density within the non-cycling progenitor cluster, suggesting that Group 3 samples exhibit a more proliferative phenotype relative to Group 4 samples (**Figure 2d**).

Quantitatively, the average proportion of each cluster within samples varied across molecular subgroups (**Figure 2e**). Group 3 tumors had the largest proportion of non-cycling progenitors and the lowest proportion of differentiating MB cells compared to other subgroups. The TAA-associated cluster comprised the highest proportion within Group 4 tumors, while the TAM- and HSR-associated clusters comprised the lowest proportion among these tumors relative to the other subgroups. Among the SHH and WNT subgroups, SHH samples had the largest proportion of voxels representing stroma, whereas the WNT tumor sample had the largest proportion of HSR-associated voxels.

Collectively, our clustering results suggest that while MB cells from distinct subgroups and cells of origin exhibit disparate genomic profiles, common biological themes, including states of differentiation, stress, and cell cycle progression, remain preserved across tumors. Furthermore, similar components of the MB-TME, including TAMs, TAAs, stroma, and tumor vasculature, are maintained across tumors. Although these samples represent a limited portion of tumors, comparing the spatial distribution and differential abundance of these MB-TME components underscores both inter- and intra-tumoral heterogeneity among molecular subgroups.

### Neighborhood enrichment analysis reveals dense regions of progenitor cells enriched in HR patients

With the annotated tumor and microenvironment clusters, we next sought to compare the differences in spatial organization between samples and clinical features using neighborhood enrichment (NE) analysis (**Figure S3**). Through the NE analysis, z-scores representing the co-localization between pairs of clusters were calculated within each sample. Increased z-scores indicate a higher proportion of one cluster within the spatial neighborhood of the index cluster. Strikingly, when comparing patients with SR versus HR disease, the neighborhood enrichment of non-cycling MB progenitors with other non-cycling MB progenitors was highly enriched in patients with HR disease. This suggests the formation of dense non-cycling MB progenitor regions in HR samples. This enrichment was significantly higher (z-score = 22.98) compared to other enriched neighborhoods within these samples and the SR cohort (**Figure 3a**). Within the SR cohort, several enriched neighborhoods were identified. Of these, TAAs were found to be enriched in the neighborhoods of differentiating, non-cycling, and HSR MB progenitors, as well as near TAMs and VWF^+^ vascular endothelium.

**Figure 3.**
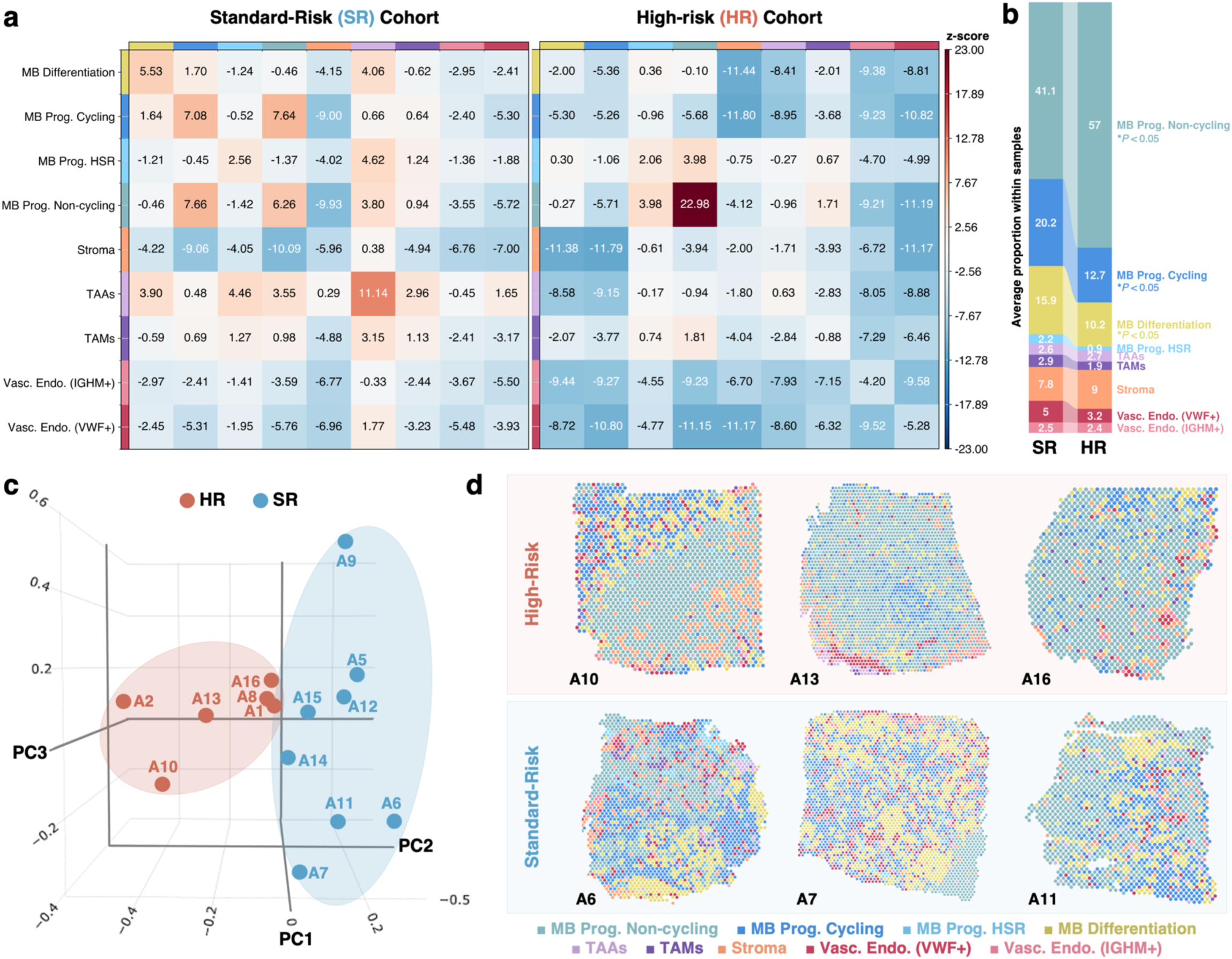
Spatial heterogeneity of high versus standard-risk patient samples showing regions of non-cycling MB progenitor cells. **(a)** Spatial neighborhood enrichment analysis of standard (*n* = 8) versus high-risk MB patient samples (*n* = 6) reveals an increase in non-cycling MB progenitor (MB Prog. Non-cycling) dense regions. Neighborhood enrichment heatmap matrices illustrate the Z-scores corresponding to the increased (*red*) or decreased (*blue*) localization of voxels from another cluster (*x-axis*) within the spatial neighborhood of voxels from the index cluster (*y-axis*). **(b)** Differential abundance of clusters between the high-risk (HR) versus stand-risk cohorts further shows a significant difference in cycling, non-cycling, and differentiating progenitors between the groups. **(c)** Principal component analysis on the per-sample spatial neighborhood enrichment from **(a)** and composition **(b)** of the samples illustrates a distinction based on risk category. **(d)** Representative spatial plots of samples from the HR and SR cohorts. Voxels are colored based on their cluster annotation labels.

Comparing the overall differential abundance of these clusters in HR versus SR disease samples, TAAs did not appear significantly enriched in SR samples (**Figure 3b**). However, their spatial localization near MB cells, TAMs, and VWF^+^ tumor endothelium suggests potential crosstalk within the MB-TME, providing insights not captured by differential abundance testing common in scRNA-seq approaches. Specifically, the secretory phenotype of astrocytes is known to play a role in shaping pathogenic phenotypes in resident brain cell populations, such as the polarization of microglia to an M2 phonotype and dysregulation of endothelial cells in the blood-brain barrier (BBB)^26,27^. Supporting our neighborhood enrichment analysis, the proportion of non-cycling MB progenitors was also significantly enriched in HR patient samples, increasing by 1.4 folds (*P* < 0.05).

Given these observed differences in the composition and spatial organization within HR patient tumors, we performed principal component analysis (PCA) based on the neighborhood enrichment and proportion of clusters present in each sample. The PCA results illustrate that samples from HR and SR patients cluster distinctly based on the variance of their spatial organization and composition (**Figure 3c**). These findings are further exemplified when examining the spatial plots of HR patient samples relative to SR patient samples (**Figure 3d**). Dense regions of non-cycling progenitors are observed across HR patient tumor samples, while SR patient tumors demonstrate a higher degree of heterogeneity (**Figure 3d**). This heterogeneity is also evident in the PCA plot, evidenced by the high variance of SR patient samples across the first three principal components, in contrast to HR patient samples.

Furthermore, the HR patient samples, A1, A8, and A16 appear to have more infiltration of other populations within their dense non-cycling progenitors regions, while SR patient samples A14 and A15 (both derived from infant SHH patients with ND type histology) appear to have developing regions of non-cycling progenitors (**Figure S1**), illustrating a continuum of non-cycling progenitor regions and suggesting signs of future disease progression.

The spatial NE analysis depicts the existence of homogenous non-cycling progenitor islands that might be associated with the progression of disease and should be further evaluated as disease progression biomarkers in a larger cohort.

### HR-associated progenitors exhibit a quiescent phenotype associated with therapeutic resistance

Following the identification of non-cycling progenitor regions, we hypothesized that these non-cycling progenitors correspond to quiescent phenotype implicated in therapy resistance. To test this hypothesis, we evaluated the expression of canonical G0 markers, associated with cell cycle arrest and quiescence, among the MB clusters^28^. Supporting our hypothesis, the non-cycling progenitor cluster demonstrated a significant upregulation of G0 markers relative to the other MB cell clusters (**Figure 4a**). Furthermore, mapping the pan-cancer G0 signature developed by Wiecek *et al.* across the MB cell clusters illustrated an enrichment of this signature in the non-cycling MB progenitor cluster (**Figure 4b**). Comparing the enrichment of this signature between SR and HR patient samples revealed a non-significant difference (**Figure S4**), indicating that the quiescence phenotype is not exclusive to HR tumors. Rather, HR tumors display an increased proportion of these quiescent progenitors in densely packed regions, as illustrated in **Figure 3**.

**Figure 4.**
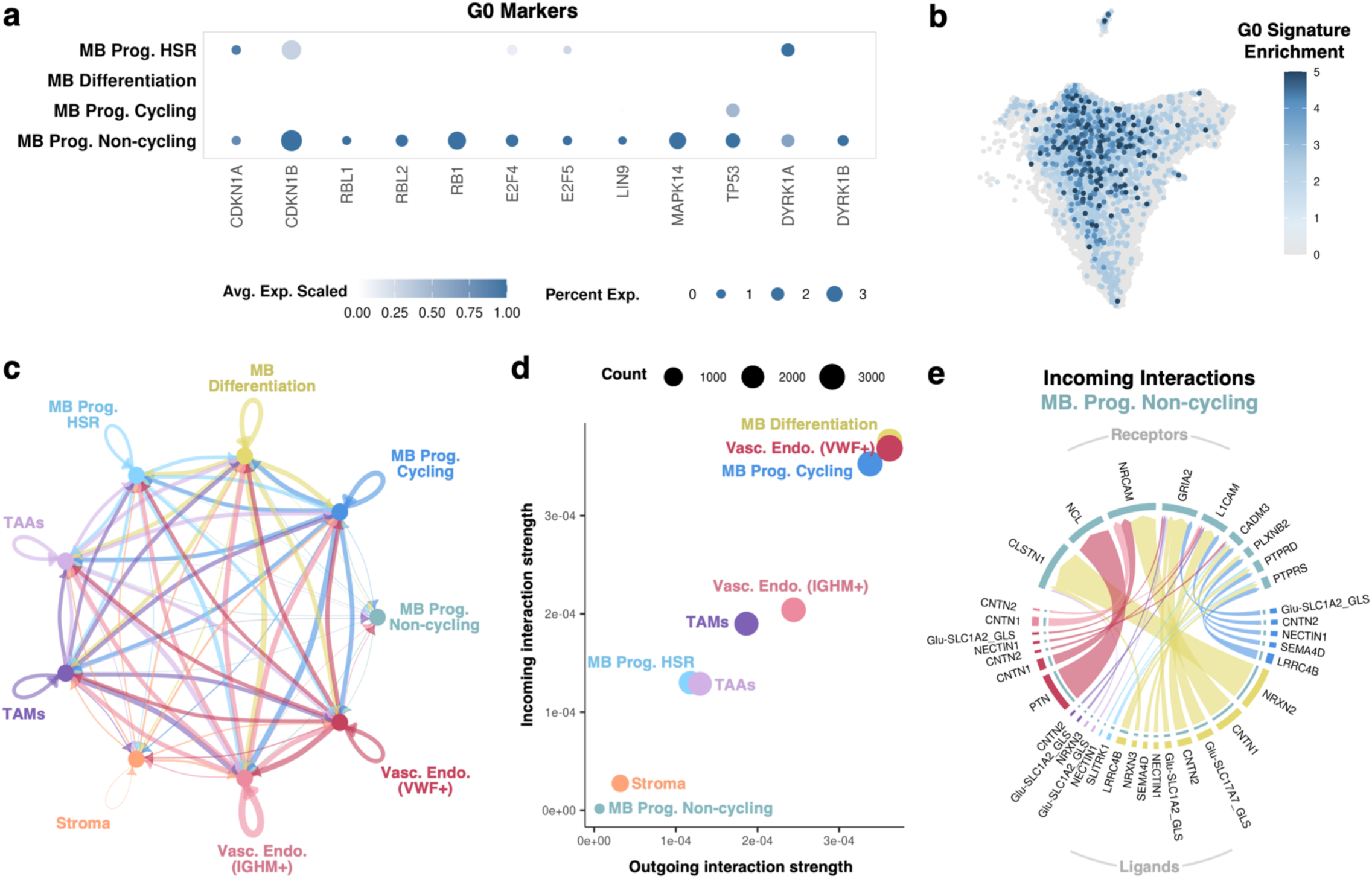
MB progenitor non-cycling cluster exhibits hallmarks of quiescence. **(a)** Dot plots of canonical markers of cellular senescence corresponding to a G0 cell state. The shade of each dot corresponds to the average expression of the gene among voxels of the cluster, while the size of each dot corresponds to the percentage of voxels expressing the gene. **(b)** Feature plot showing enrichment of the senescent G0 signature across MB cell clusters. **(c-e)** Intercellular communication analysis was performed on samples at diagnosis. Colors correspond to the respective clusters. **(c)** Chord diagram illustrating the number of spatially dependent cellular interactions between labeled voxels, highlighting limited cellular signaling by quiescent MB progenitors (MB Prog. Non-cycling). The width of the chords corresponds to the strength of the interactions. **(d)** Scatter plot displaying the strength of outgoing (ligand) and incoming (receptor) signals for each cluster. **(e)** Chord diagram of the incoming signaling interactions to the non-cycling progenitor cluster signaling illustrates the ligands expressed by each cell type in close proximity to the non-cycling MB progenitors (*bottom*) and their respective receptor expressed on non-cycling MB progenitors (*top*). The width of each chord corresponds to the strength of signaling interaction.

We next questioned whether another cell type within MB-TME may be driving this quiescent phenotype in MB cells. To examine this, we performed spatially dependent intercellular communication analysis (see Methods), comparing the co-expression of ligand-receptor pairs in close spatial proximity across samples. Consistent with a quiescent phenotype, these non-cycling progenitors predominantly received weak incoming signals from neighboring cells with no significant outgoing signaling interactions, including autocrine signaling (**Figure 4c**). In contrast, VWF^+^ vascular endothelium, differentiating MB cells, and cycling MB progenitors exhibited high degrees of intercellular signaling interactions within the TME (**Figure 4d**).

While the overall signaling interactions with the non-cycling progenitors were weak, potentially due to the spatial isolation of these cells, we examined the incoming signals (ligands) interacting with receptors expressed on the non-cycling progenitors, identifying several interesting interactions (**Figure 4e**). First, the multifunctional protein nucleolin (NCL) appears to be stimulated by the growth factor pleiotrophin (PTN) produced by VWF^+^ vascular endothelium. Notably, increased nucleolin expression has been implicated in chemotherapy resistance for several cancers, including resistance to platinum-based therapies^29,30^. Furthermore, putative roles for pleiotrophin–nucleolin interactions in angiogenesis and metastasis have been reported^31,32^. Additionally, these non-cycling progenitors appear stimulated by pleiotrophin and contactin 1 (CNTN1) binding to the neuronal cell adhesion molecule (NrCAM) receptor. Contactin 1 exhibits pleiotropic functions across many cancers, facilitating tumor progression, therapy resistance, and metastasis^33,34^. Lastly, neuronexin 2 (NRXN2) binding to calsyntenin 1 (CLSTN1) also emerged as a significant signaling interaction in non-cycling progenitors. The cadherin-family transmembrane protein calsyntenin 1 is involved in axon development and vesicle transport; however, its function in cancer remains unexplored^35^. Collectively, the signaling pathways involved in non-cycling progenitors appear to be linked to migration and therapy resistance, warranting further investigations into their role in MB.

### Intercellular communication analysis reveals metastasis-associated signaling patterns in HR disease

To compare the TME of HR tumor samples to that of SR tumor samples, we extended our intercellular communication analysis to identify significant differentially enriched signaling patterns between HR and SR tumors (**Figure 5a**). In HR tumors, increased outgoing (ligand) and incoming (receptor) signaling was observed among VWF+ endothelial cell and TAA clusters, predominantly mediated by claudin (CLDN), tubby-like protein (TULP), and laminin interactions. In contrast, SR tumor samples primarily exhibited increased signaling mediated by laminins, neuregulins (NRG), and contactins (CNTN) in the differentiating MB cell cluster.

**Figure 5.**
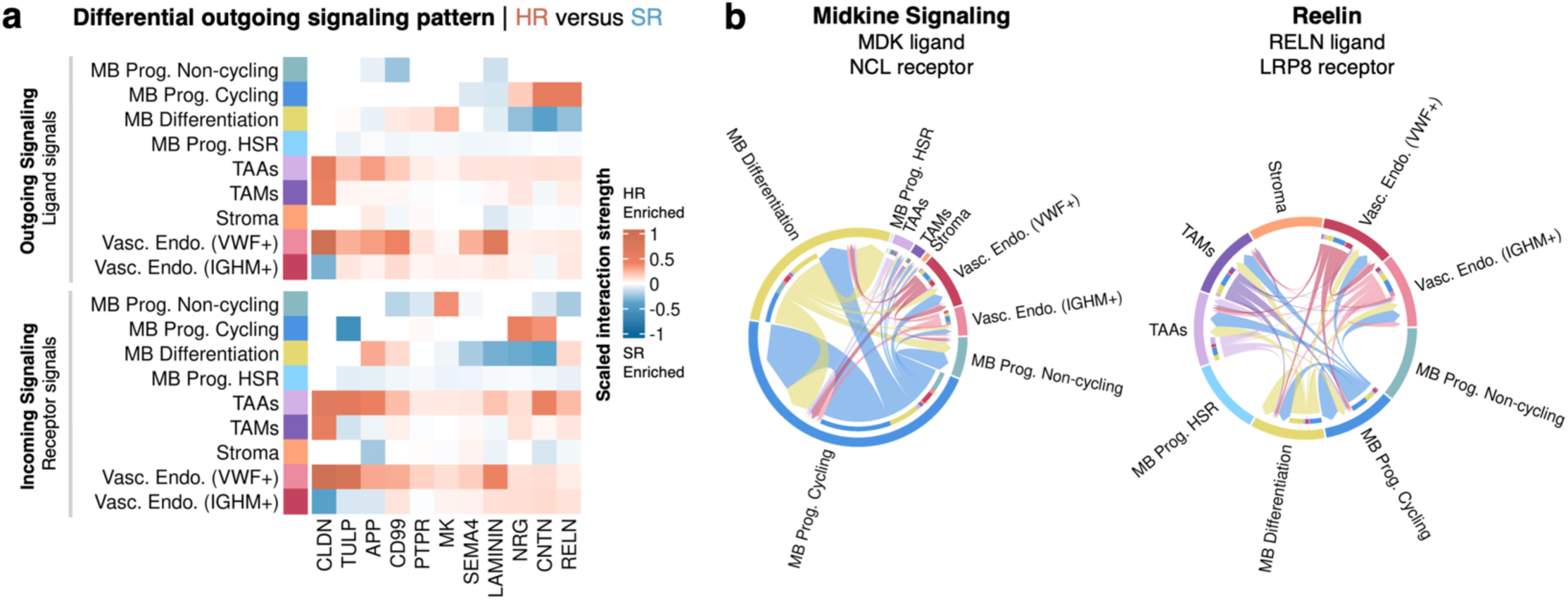
Spatially dependent intercellular communication analysis reveals metastasis-associated signaling patterns in HR disease. **(a)** Heatmap depicting the differential incoming and outgoing signaling patterns between high-risk (HR, *red*) and standard-risk (SR, *blue*) cohorts. Notably, intercellular signaling mediated by TAAs and VWF^+^ vascular endothelium is increased in HR relative SR samples. Furthermore, non-cycling MB progenitors receive increased stimulation by the pro-tumoral midkine (MK) molecule in HR samples, while cycling MB progenitors exhibit increased reelin (RELN) production. **(b)** Chord diagrams showing midkine and reelin signaling pathways associated with MB disease progression and metastasis in HR patient samples.

Among the differentially enriched signaling patterns, midkine (MK) and reelin (RELN) mediated signaling stood out, as these signaling pathways were enriched in the non-cycling and cycling MB progenitors of HR tumor samples, respectively. Closer examination of these signaling patterns in HR tumors revealed that MK was primarily secreted by differentiating and cycling MB cells and bound to the NCL receptor expressed on non-cycling MB progenitors (**Figure 5b**). Conversely, RLN was also secreted by differentiating and cycling MB cells but interacted with LDL receptor related protein 8 (LRP8) receptors expressed on glial and endothelial cells (**Figure 5b**).

Importantly, midkine is a well-studied growth factor involved across several tumor-promoting pathways^36^. Its increased secretion by cycling and differentiating MB cells, received by the NCL receptor on non-cycling MB progenitors, may promote the reactivation of these quiescent cells, promoting relapse and metastasis. Reelin also plays a well-documented role in tumor progression, including a recent report of its function in MB metastasis^37^. Its primary role in cell migration suggests that reelin-mediated signaling amongst glial, endothelial, and MB cells in the TME of HR tumor samples may promote tumor growth in HR patients by stimulating angiogenesis and tumor proliferation.

### HR-associated progenitors exhibit a quiescent phenotype associated with therapeutic resistance

Given that our dataset uniquely included two patient-matched samples from diagnosis and relapse, we compared these matched samples to identify differences in the cellular and spatial architecture before and after relapse (**Figure 6**). The first matched pair (A1 diagnosis, A3 relapse) derives from a patient (Patient 1) who presented with a Group 4 tumor with classical histology, isochromosome 17q, and M2 stage metastasis at diagnosis. The second matched pair (A2 diagnosis, A4 relapse) derives from a patient (Patient 2) who presented with an SHH tumor with *MYCN, GLI2*, *PPM1D*, and *TERT* alterations and LC/A histology. Spatial comparison of diagnosis and relapse tumor samples shows uniform regions of non-cycling progenitors, consistent with their HR designation in diagnostics samples (**Figure 6a**). Following relapse after frontline therapy, tumors appear more heterogeneous, with constituents of the tumor microenvironment more interspersed with less apparent spatial separation between malignant and stromal cells.

**Figure 6.**
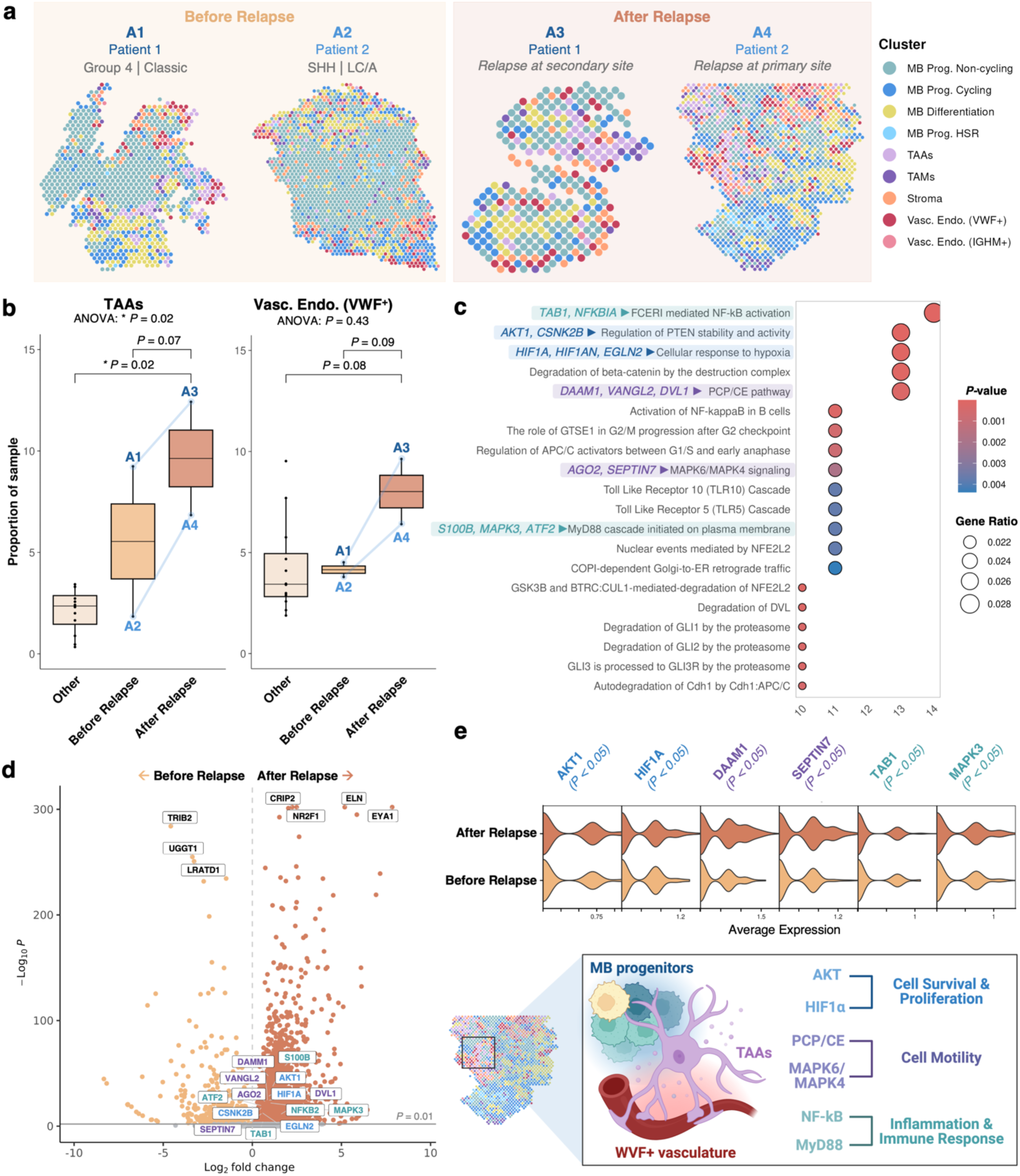
Dysregulation in TAAs and vasculature were observed in matched diagnosis-relapse MB samples. **(a)** Spatial plots of samples before and after relapse. **(b)** Box plots illustrating the proportion of TAAs and VWF^+^ vascular endothelium in relapsed tumor samples relative to matched samples before relapse (at diagnosis) and samples without documented relapse (other). **(c)** Pathway enrichment (Reactome database) based on genes upregulated between TAAs and VWF^+^ endothelium in close proximity within relapsed mem. **(d)** Volcano plot displaying the differentially expressed genes before and after relapse. The top genes ranked by adjusted *P*-value before and after relapse are highlighted in black, while key genes from enriched pathways in **c** are highlighted in blue, purple, and green, corresponding to pathways related to cell survival and proliferation, cell motility, and inflammation and immune response, respectively. **(e)** Violin plots displaying the average expression of differentially expressed genes (*P* < 0.05) involved in key pathways related to cell survival and proliferation (blue), cell motility (purple), inflammation and immune response (green), among TAA and VWF^+^ vascular endothelium clusters before and after relapse.

Comparison of the patient-matched samples illustrates increased proportions of TAA and VWF^+^ vascular endothelial cell clusters at relapse compared to diagnosis and unpaired samples (*P* < 0.05), suggesting that these populations may play a role in relapse (**Figure 6b**). Considering that a key function of TAAs is the modulation of the BBB, we next performed niche-dependent differential gene expression analysis (see Methods) to elucidate how the transcriptomic profiles of VWF+ vascular endothelium and TAAs are altered when these cell types are in close proximity. A total of 2,293 significant differentially expressed genes (*P* < 0.05) were identified in neighboring TAAs and VWF^+^ vascular endothelium relative to their nonadjacent counterparts, pointing to a spatially dependent (niche) phenotype for these cell types. Further pathway enrichment analysis on these differentially expressed niche genes identified several significantly enriched pathways (*P* < 0.05) (**Figure 6c**). Namely, the upregulation of the AKT serine/threonine kinase 1 (*AKT1*) and casein kinase 2 (CK2, *CSNK2B*) in the PTEN pathway promoting cell survival and growth, in addition to hypoxic response genes hypoxia-inducible factor 1-alpha (*HIF1A*) and inhibitor (*HIF1AN*) with egl-9 family hypoxia-inducible factor 2 (*EGLN2*). Furthermore, enrichment of the planar cell polarity (PCP) pathway involved in non-canonical WNT signaling and the MAP kinase 4/6 signaling pathways, both with central functions in cell migration and invasion, were observed. Additionally, pro-inflammatory and immune response pathways mediated by nuclear factor kappa-light-chain-enhancer of activated B cells (NF-kB) and myeloid differentiation primary response 88 (MyD88) signaling were also enriched. Furthermore, the key genes identified in these enriched pathways were among the overall differentially expressed genes following relapse (**Figure 6d-e**). Considering that each spatial voxel is non-homogenous and contains approximately 1–10 cells, voxels within the TAA and VWF^+^ vascular endothelium clusters also contain other cell types, including potential pro-metastatic cells that may intravasate through a dysregulated blood-brain barrier (**Figure 6e**). Thus, taken together, these results may indicate that pro-metastatic niches are formed in regions of TAAs and VWF^+^ vascular endothelium, facilitating relapse.

## Discussion

The medulloblastoma TME is known to influence tumor progression and relapse, thus targeting components of the TME has emerged as a novel therapeutic strategy^9^. Prior studies, including single-cell transcriptomic analyses, have described the cellular composition of the MB-TME^10,11^, but in these studies, the spatial association of cell types is lost. Elucidating the spatial architecture of the TME can provide critical insights into mechanisms of disease progression, by revealing localized niches, spatially dependent patterns of cellular signaling, and the co-localization of specific cell types. Through spatial sequencing of human medulloblastoma tumor tissue, we characterize both the cellular composition and, importantly, the spatial organization of the medulloblastoma TME.

The cell types identified through our clustering analysis revealed clusters corresponding to TAMs, TAAs, vasculature, and stromal components in addition to malignant cells, aligning with previous studies of the MB-TME. TAMs in CNS malignancies represent a diverse population consisting of both resident microglia and monocyte-derived macrophages with phenotypic plasticity^38^. The TAM cluster of our dataset, comprised of both microglia and monocyte-derived macrophages, highly expressed *SPP1*, an emerging biomarker of protumoral TAMs and a contributor to chemoresistance in solid tumors^21,22,39^. The role of SPP1^+^ TAMs in MB and the mechanisms by which they may promote disease progression remain to be described; however, their enrichment in relapsed samples within our dataset suggests that these cells warrant future investigations.

Among malignant cells in the MB-TME, our identification of progenitor, differentiating, and cycling subpopulations aligns with MB cell subtypes previously characterized by single-cell analysis^10,11,40^. Interestingly, in addition to these previously described subtypes, we also identified a unique subpopulation of malignant cells with elevated expression of HSR proteins, namely HSP90 and HSP70 family proteins, implicated in tumor progression^23,24,25^. Moreover, through our spatial analysis, we provide novel insight into the orientation of these cell types across the TME. Specifically, by comparing correlations of spatial composition to molecular subgroups and clinical features, we highlight the intertumoral heterogeneity previously inferred through multiregional surgical biopsy^1,6,41^. Notably, we identified the presence of high-density non-cycling progenitor regions correlating with disease at high risk of relapse. These non-cycling progenitor cells express markers of quiescence and decreased intercellular signaling, supporting their role as dormant malignant cells. Dormancy confers tumor cell resistance to anti-neoplastic therapy^18^, suggesting the increased presence of non-cycling progenitors may drive future disease progression and relapse in MB, following which no curative therapy is available. Collectively, our findings emphasize the heterogeneity of the MB-TME and further suggest the potential limitations of frontline DNA-alkylating agents that may not target the entire malignant cell population.

To further probe potential intercellular signaling patterns promoting the observed non-cycling progenitor phenotype, we examined the spatially dependent cellular communication patterns between cells in the MB-TME, identifying signaling patterns related to precursor neuron function and tumor progression. Incoming signals to these non-cycling progenitors were primarily characterized by pleiotrophin- and contactin-1-mediated signaling. Pleiotrophin, facilitated intracellularly by the cell surface receptor nucleolin, is a marker of neural stem cells and involved in neuron maturation^42^. Pleiotrophin-nucleolin interactions have also been implicated in chemotherapy resistance, cancer growth, and metastasis^29,30,32^. Contactin-1 is a cell adhesion molecule that has emerged as an oncogenic protein promoting cancer progression and metastasis^33,34^. In aggregate, these findings further support the role of non-cycling progenitor cells as dormant cancer cells with the capacity to drive future disease recurrence.

Matched samples from diagnosis and relapse for two patients are present within our dataset, providing a unique opportunity to characterize changes to the MB-TME following treatment and relapse. The two samples obtained at relapse notably differed from matched samples at diagnosis, with an increase in intracellular tumor heterogeneity, as well as an increase in the proportion of TAA and vascular endothelium-related clusters, suggesting a potential role in disease progression^43^. This increased proportion of vascular endothelium expression may represent chemoradiation-induced angiogenesis or a response to hypoxia during tumor progression and metastasis. Furthermore, recent studies have identified a putative pro-tumoral role for TAAs in the MB-TME, promoting MB metastasis^15,16,17^. Niche differential gene expression for regions of co-localizing TAAs and VWF^+^ vascular endothelium revealed genes associated with protumoral pathways known to be activated in MB, including those related to cell survival and proliferation (AKT, HIF1A), cell motility (PCP/PE, MAPK6/4), and inflammation and immune response (MyD88, NF-kB). Given the role of TAAs and vascular endothelium in maintaining the BBB, these results may suggest that protumoral TAAs dysregulate the BBB through endothelial and MB cell activation, facilitating relapse. Additional investigation using spatial sequencing of an expanded cohort of matched MB tumor samples before and after treatment would be necessary to confirm these findings and further address these questions.

While our work presents a significant milestone in elucidating the MB-TME, it has several limitations. First, while sequencing-based spatial transcriptomic approaches enable unbiased analysis compared to probe-based methods, the Visium platform is considered a low-resolution spatial transcriptomic approach, wherein each voxel contains approximately 1–10 cells. While spatial information is preserved, compared to single-cell transcriptomics, gene expression is averaged among the cell types comprising a single voxel. Therefore, careful consideration must be taken when interpreting results, noting that expression profiles correspond to sets of cells. Next, while TAMs, the most common immune constituent of MB-TME, were identified, rarer immune subtypes, such as TILs, were not well-represented across samples, likely due to the limited sample area. Future investigations integrating single-cell RNA-seq datasets may enable deconvolution of the presented spatial dataset, unveiling the spatial localization of rare cell types and subtypes identified through single-cell analysis. Thus, the spatial dataset presented herein provides a foundation for future studies of the MB-TME integrating these approaches. Finally, the dataset was obtained from a patient cohort selected based on the availability of high-quality tissue at the time of initial resection for spatial sequencing. Within our study, 13 of 14 patients are male, which overrepresents male-specific differences in tumor microenvironment. Epidemiologically, MB is overall more common in male than female patients, with a 1.8:1 male-to-female ratio^6^. Whether there are sex-dependent differences in medulloblastoma TME is not well understood. However, sex differences are noted in molecular and methylation patterns of disease, with estrogen receptor β signaling hypothesized to have a tumor-suppressive role in medulloblastomas^44–46^.

Spatial transcriptomics represents a novel strategy for characterizing the cellular composition and spatial architecture of the MB-TME. This work underscores the ability of spatial transcriptomics to capture distinct patterns of cellular organization associated with clinical features, which are lost in traditional single-cell RNA-seq approaches. Most notably, regions of non-cycling progenitor cells may be predictive of therapy response and may prove a novel biomarker for HR disease. To address this, drug-induced cell cycle modulation may be explored in the future as a means of targeted therapy in the MB-TME. Additionally, the identification of co-localizing TAAs and dysregulated vasculature following relapse may present an opportunity for exploring adjuvant therapies that target TAAs to sensitize tumors to upfront chemotherapy, decreasing relapse and mortality from disease.

## Methods

### Study Population

This study was approved by the Emory University Institutional Review Board (IRB), Atlanta, GA. Signed informed consents were obtained from all participants or legal guardians to permit the use of biological material in accordance with IRB approval. Study participants were not compensated for their participation. Spatial sequencing was performed on tissue from 14 patients treated at Aflac Cancer and Blood Disorder Center of Children’s Healthcare of Atlanta. Samples were obtained at diagnostic resection from all patients. Additionally, 2 patient-matched samples were obtained from biopsy at relapse. All sample slides were visually inspected by neuropathology and high-quality viable tumor regions of interest were circled for spatial transcriptomics.

### Methylation profiling

Genome-wide DNA methylation was performed at the New York University (NYU) Langone Health Molecular Pathology Clinical Laboratory Improvement Amendments (CLIA)-certified laboratory using the Illumina Human Methylation EPIC array as described previously^47^ and analyzed using the Heidelberg (DKFZ)-developed and NYU-clinically validated DNA methylation classifier^48^. All cases scored with a calibrated score > 0.9, which is considered positive. Copy number plots were generated using *conumee* package and reviewed visually to correlate with other molecular analyses.

### Risk stratification

First biopsy samples from 14 unique patients within our cohort were stratified into high-risk (*n* = 6) or standard-risk (*n* = 8) following surgical resection according to Children’s Oncology Group risk definitions^7^. Infants and young children who received radiation-sparing regimens with high-intensity chemotherapy were assigned high-risk if presenting with metastatic disease (M+ Chang staging) or standard risk if presenting with M0 disease and no other high-risk molecular features (e.g. *MYC*-, *MYCN*-amplification). Large-cell anaplasia as the sole determinant of higher risk is currently of undetermined clinical significance and did not independently constitute high-risk designation. The detailed patient characteristics have been included in **Supplemental Material 1.**

### Visium tissue permeabilization optimization, gene expression library construction, and sequencing

The Visium spatial gene expression platform (10x Genomics) enables the analysis of RNA levels in Formalin Fixed Paraffin Embedded (FFPE) tissue sections by utilizing probes designed to target the entirety of the transcriptome. Tissue sections were cut by Microtome (cat#23-900-672, Thermo Fisher Scientific) to a thickness of 5 µm, placed in a warm water bath (42 °C) for 1 minute, then transferred to the capture area (CG000408 Rev E, 10x Genomics). Deparaffinization, hematoxylin and eosin staining (H&E), and decrosslinking were performed based on the manufacturer protocol (CG000409, Rev D, 10x Genomics). Brightfield images of the H&E stained tissues were taken using a 40X objective on NanoZoomer 2.0 HT. Library constriction was performed by following manufacturer protocol (CG000407 Rev E, 10x Genomics). The gene expression libraries were quantified using a Qubit fluorometer (Thermo Fisher Scientific) and checked for quality using DNA HS bioanalyzer chips (Agilent). Sequencing depth was calculated based on the percent capture area covered by the tissue, and the 10x Genomics recommended sequencing parameters were used to run on the NovaSeq 6000 with S4 PE 100 kits (Illumina).

### Spatial data processing and analysis

FASTQ files for each sample were aligned to the human GRCh38 genome using Space Ranger (v2.0.0, 10x Genomics). Microscopy images of the H&E stained samples were aligned to the spatial voxels through the Space Ranger pipeline. Noncontiguous portions of the tissue present in the capture area were manually annotated and removed from downstream analysis. Samples ranged in size from 1,000–3,000 voxels with a median of 2,524 transcripts captured per sample and 16,238 unique genes represented. Spatial plots of each sample were generated using Seurat. Spatial plots in the main text illustrate representative samples, while additional spatial plots are provided in Figure S1. The metadata corresponding to each sample was visualized using an upset plot from the UpSetR package^49^.

The samples were normalized using the *SCTransform* function and then integrated using integration anchors-based batch correction via the Seurat R package (v4.4.0)^50^. Principal component analysis (PCA) was then performed on the integrated spatial and single-nuclei datasets using all non-ribosomal features, followed by construction of a K-nearest neighbors graph (using cosine similarity) and Louvain clustering, then non-linear dimensional reduction via uniform manifold approximation and projection (UMAP). The resolution (*n* = 0.8) and 30 principal components were used for clustering selected based on the Clustree R package (v0.5.0)^51^.

Differential gene expression analysis was performed via Seurat using the *FindAllMarkers* function, comparing expression between clusters without spatial consideration with a log fold-change threshold of 0.1. Significant differentially expressed genes were determined based on the Wilcoxon rank sum test with Bonferroni multiple test correction (*P* < 0.05). Gene expression dot and violin plots displaying differentially expressed genes and canonical markers were generated using Seurat. Volcano plots displaying differentially expressed genes were generated using EnhancedVolcano (v1.20.0). Differential abundance testing between multiple groups (ANOVA) was performed using the Speckle R package via the *propeller* function with arc sine scalar transformation, robust empirical Bayes shrinkage of the variance, and Benjamini and Hochberg (BH) correction^52^. For differential abundance testing between unpaired samples, a Wilcoxon rank-sum test was performed with BH correction, while a paired t-test was used to compare samples before and after relapse. Box plots and bar plots of the cluster proportions were generated using ggplot2 (v3.5.1).

### G0 Cell Identification

To map the enrichment of the pan-cancer G0 cell signature reported by Wiecek *et al.*, the module scoring was performed via Seurat using the *AddModuleScore* function with default parameters^53^. The G0 signature contained the following 27 genes: *CFLAR, CALCOCO1*, *YPEL3*, *CST3*, *SERINC1*, *CLIP4*, *PCYOX1*, *TMEM59*, *RGS2*, *YPEL5*, *CD63*, *KIAA1109*, *CDH13*, *GSN*, *MR1*, *CYB5R1*, *AZGP1*, *ZFYVE1*, *DMXL1*, *EPS8L2*, *PTTG1IP*, *MIR22HG*, *PSAP*, *GOLGA8B*, *NEAT1*, *TXNIP*, *MTRNR2L12*.

### Neighborhood enrichment analysis

Neighborhood enrichment analysis was performed on samples from diagnosis employing the voxel labels derived from clustering. Using the *gr.spatial_neighbors* function via the Squidpy python package (v.1.4.1)^54^, a z-score was generated for each cluster pair based on the enrichment of the respective cluster within the neighborhood (60 total voxels) of voxels of the index cluster, where a higher score represents closer proximity. To generate the enrichment plot, the connectivity matrix was calculated using the *gr.nhood_enrichment* function and plotted using the *pl.nhood_enrichment* function.

### Principal component analysis

Min-max normalization was applied to the values of the per-sample neighborhood enrichment connectivity matrices and cluster proportions, generating normalized values between −1 and 1 across samples (at diagnosis only) for each cluster and paired cluster neighborhood enrichment z-score. Principle component analysis was performed using a matrix of these values as input to the *prcomp* function in the base R stats package (v4.3.0) with data centering. A 3D plot of the first three principal components was generated using the Plotly R package (v.4.10.4) to visualize the results.

### Niche-dependent gene expression

Niche-dependent differential gene expression analysis was performed using the NicheDE R package (v0.0.0.900). The NicheDE package examines differential expression (DE) across a tissue sample by calculating how the gene expression of an index cell type changes when in close spatial proximity to another cell type. The composition of cell types surrounding an index cell is termed its “effective niche” and represents the biological niche which can influence the expression of cell type through intercellular signaling. For example, the gene expression profile of a vascular endothelial cell may change when in close proximity to TAAs secreting matrix metalloproteases. For our Visium spatial dataset, the cell type of a voxel is represented by the cluster annotation label of the voxel. For example, all voxels of the TAA cluster are labeled as TAA cells; however, TAAs are the predominant cell type of the voxel, and additional cells may be present within these voxels.

To perform the niche DE analysis, a nicheDE object was created for each tumor sample using the spatial count matrix generated by Seurat. Sigma values (corresponding varying sizes of the effective niche) of 25, 50, 75, and 100 were assessed, with a value of 75 selected for downstream analyses. The resulting sample-wise objects were merged, and the effective niches (composition of cell types in surrounding voxels) were calculated using the *CalculateEffectiveNiche* function with a cutoff value of 0.05. The niche-dependent DE for each cell type was then performed using a minimum total gene expression cutoff (C) of 50 with a minimum of 5 spots containing the index cell type (M) using the *niche_DE* function. To determine how the gene expression profiles of TAAs change when in the effective niche of (i.e., close proximity to) VWF^+^ vascular endothelium, the niche DE genes were then calculated using voxels from the TAA cluster within the effective niche of voxels from the VWF^+^ vascular endothelium cluster. Similarly, the niche DE genes were calculated for VWF^+^ vascular endothelium voxels within the effective niche of TAA voxels. These genes were then filtered to only include significant genes (*P* > 0.05) with ribosomal genes regressed.

### Pathway enrichment analysis

Pathway enrichment analyses were performed using the ClusterProfiler R package (v4.10.1) and the Reactome database. The differentially enriched gene sets resulting from the niche-dependent gene expression analysis, were converted to Entrez Gene IDs and used as input for the *compareCluster* function. Significantly enriched pathways were calculated using the *enrichPathway* function, which uses the hypergeometric distribution to calculate *P*-values with Bonferroni multiple test correction (*P* < 0.05). Pathways specific to infectious diseases were manually filtered out. To eliminate redundancy among significantly enriched pathways, the pathway with the lowest enrichment q-value was selected from significant pathways containing identical gene sets. Dot plots of enriched pathways were generated via the *dotplot* function in ClusterProfiler.

### Intercellular communication analysis

Intercellular communication analysis was performed using the CellChat R package (v2.1.2)^55^. Communication probabilities were calculated using the *computeCommunProb* function with a distance constraint and contact range of 100, a minimum of five interacting cell pairs, and a scale factor of 1. Analysis was performed on all 16 samples first as a single CellChat object, then samples from diagnosis were subset into the HR and SR groups and analyzed separately. The HR and SR communication analyses were combined via the *mergeCellChat* function to compare the differences in their intercellular communication networks between groups (HR vs SR) using previously described methods^55^. To identify shared communication pathways between groups, the intersection of significant (*P* < 0.05) signaling pathways between the groups was used. The differential heatmap of interaction strength was generated by subtracting the overall interaction matrix of the HR patient samples relative to SR patient samples, removing significant signaling pathways with <0.01 change in signaling strength, and visualized using the ComplexHeatmap R package (v.2.18.0).

## Data availability

The spatial sequencing data that support the findings of this study are available via the Gene Expression Omnibus database under accession number GSE000000 (pending).

## Acknowledgments

This work was funded by the CURE Childhood Cancer Foundation. DNA methylation analysis was in part supported by National Institutes of Health (5R01NS122987 to M.S.) and the Friedberg Charitable Foundation (to M.S.).

The content is solely the authors’ responsibility and does not necessarily represent the official views of the National Institutes of Health. Figure illustrations were made with BioRender.com.

## Article Information

### Contributions

Conception and design: F.C., T.M., M.K.B. Development of the methodology: M.B., M.S., C.S. Acquisition of data: M.B., M.S., C.S., M.S. Analysis and interpretation of data (bioinformatic analysis): M.M., F.C. Writing, review, and/or revision of the manuscript: F.C., M.E.M., T.M., M.K.B.

### Corresponding authors

Correspondence to Tobey MacDonald or Manoj Bhasin.

## Ethics declarations

### Competing interests

M.S. is scientific advisor and shareholder of Heidelberg Epignostix and Halo Dx, and a scientific advisor of Arima Genomics, and InnoSIGN, and received research funding from Lilly USA. The other authors declare no competing interests.

## Notes

### Competing Interest Statement

The authors have declared no competing interest.

