## Supplemental Material 2 for "Medulloblastoma Spatial Transcriptomics Reveals Tumor Microenvironment Heterogeneity with High-Density Progenitor Cell Regions Correlating with High-Risk Disease"

### Table of Contents

**Supplemental Table**

**Table S1.** Table listing the gene program signatures used for annotations………………………..……………… 3

**Supplemental Figures**

**Figure S1.** Spatial plots illustrating the spatial distribution of clusters across individual samples………………. 4

**Figure S2.** Expression of *SPP1* corresponding to clinical features……………..…………………………………..…. 5

**Figure S3.** Neighborhood enrichment analysis z-score matrices by sample and clinical feature…….…………… 6

**Figure S4.** Enrichment of G0 signature in HR versus SR samples……………………......…………………….. 7

**Table S1.** Table listing the gene program signatures derived from Hovestadt *et al.* used for annotations.

| Program | Genes |
|---|---|
| **Cycling Program** | *TOP2A, PBK, UBE2C, ZWINT, BIRC5, CDK1, NUSAP1, TACC3, NUF2, TYMS, MELK, TPX2, AURKB, ASF1B, CDCA5, BUB1B, CENPF, KIF23, TROAP, NDC80, RRM2, KIF2C, FAM64A, KIAA0101, CDCA8, UBE2T, CCNA2, SPAG5, MXD3, CDKN3, MLF1IP, CDC20, CCNB2, CENPM, TK1, CDCA3, PTTG1, MAD2L1, KIF11, CENPN, PRC1, NMU, CENPK, UHRF1, NCAPD2, FANCI, SMC4, DEPDC1B, OIP5, DHFR, CCNB1, HMGB2, NCAPG, FAM83D, SGOL1, RAD51AP1, ECT2, FBXO5, ORC6, CENPH, NEK2, SMC2, RNASEH2A, LRR1, LMNB2, CDC6, DTYMK, CHEK2, KIAA1524, HIST1H3I, CDCA4, GMNN, CENPW, CKS2, KIF22, PLK1, MIS18BP1, HIST1H4C, RFC3, CKS1B, TUBA1B, AURKA, NCAPG2, HMGB1, MNS1, DSN1, CENPL, DEK, HELLS, PCNA, KPNA2, LIG1, LMNB1, TRIP13, MCM3, GINS2, TMPO, H2AFZ, CKAP2, POLA2, DTL, RACGAP1, LDHA, MZT1, KIF18B, SAE1, H2AFY, EZH2, FEN1, VRK1, TIMELESS, TUBB6, XRCC2, DNAJC9, PSRC1, TUBB4B, FANCA, DNMT1, IDH2, PRR11, DDX39A, RAD21, TMEM106C, PHGDH, MASTL, CMC2, NUCKS1, C5orf34, RRM1, KIFC1, PKMYT1, HJURP, CKAP2L, ESCO2, CLSPN, FANCD2, EXO1, BRCA1, ATAD2, WDR76, SKA3, CDC45, TTK, ATAD5, KNTC1, RAD51, HMGN2, PSMC3IP, HIST1H1B, HAUS8* |
| **Progenitor Program** | *IFT57, GLT8D2, RPL8, RPSA, RPLP0, RPS3, PLXDC1, C17orf76-AS1, RPL3, ASB13, RPS7, NME2, PAPSS1, ADAMTSL1, GNB2L1, VSTM2A, MFAP4, RPL6, DKK2, RPL4, ASIC2, PRDX1, PDE11A, CA4, RPS8, EEF1G, MEIS1, PGM5-AS1, CACNA2D4, HK2, RPS6, RPS3A, RPL7A, RPL29, NET1, RPL7, RPL13, SLC26A7, COL16A1, LY6E, SEC61A1, USH2A, LGALS3BP, RPL5, PGM5, RPL18A, RPLP1, COL5A1, NELL2, HM13, TMEM51, PCSK9, RPS17, CRABP2, PABPC1, MDK, NPM1, KCNH5, RPL19, RPL9, AMHR2, ABHD14A, MAF, EMB, AMBRA1, SPRY2, MAP2K6, RPS24, RPS14, NACA, HLA-A, TBCK, OSTC, PPIB, EPHA4, AIMP1, RPL11, P4HB, RPL35, GAS5, CDHR1, RPS2, TSPAN3, RPL13A, GPX1, RPL17, IGSF21, RPL15, NTPCR, RPN2, RPL22, RPL14, ST6GAL2, NTN5, LAMB1, RPL27, PDIA4, ZNF385C, TNC, RPL23A, SERPINF1, ALDH1A3, MGP, NRIP2, CLDN1, EXOC3L1, RPL10A, EEF1A1, RPS11, GPR78, BOC, RPS12, RPL23, RPS18, RPL12, PTCH1, RPL41, RPL32, GFRA1, PGM1, FOXS1, RPS15A, RPL26, RPL28, RPS9, RPS19, ANGPTL2, NDST3, PTCH2, ELK2AP, RPL30, RPS27A, RPL36, RPL35A, RPS25, RPL27A, CPZ, PCNT, WWTR1, RPS27, RPS20, RPS10, RPL31, RPS23, RPS16, RPS28, RPL34, EIF3E, RPL37, VSTM4, SEPP1, RPL18, RPS5, MTRNR2L1, RPS13, CSNK1E, MTRNR2L2, RPS15, SLC12A4, RPL37A, RPLP2, MTRNR2L8, RPL24, GAK, MXRA8, PRAME, SNHG8, MYC, TSPO, HLX, FTL, EIF3H, SLC1A7, LDHB, LAPTM4B, DPP7, RPS29, SNHG5, TKT, HIST1H2AC, COX7C, EPB41L4A-AS1, SDC2, BTF3, TOP1MT, DCN, PPA1, IFITM3, UQCRB, GSTP1* |
| **Neuronal Differentiation Program** | *LINC00599, STMN4, KIF5C, CADPS, THSD7A, MIAT, KIAA1107, DPYSL3, DUSP26, MAP1B, NHLH1, TACC2, DCC, CADM3, IMPG2, CASP3, TMEM59L, ATCAY, STMN2, SH2D3C, PTPRO, TPH1, PIK3R1, ABCC8, SSTR2, EBF1, SYT11, ATP1A3, LRRTM2, ST6GALNAC5, FAIM2, H3F3B, TUBB3, C11orf87, PTPRN, BTBD17, PKIA, AKAP6, KIF5A, GAP43, SYT1, SPOCK2, GNG3, ENO2, SCG2, TTC3, TAGLN3, RBFOX2, ITGB8, CCND2, TP53INP2, NTRK1, MEG3, TUBA1A, CALM1, CLASP2, RTN1, CELF4, PRPH, MAP1LC3A, PTPRS, STXBP1, OCIAD2, ZFHX4, FAM174B, STX3, LAMA5, NEUROD4, SOX4, SCARB2, PCDH9, MPP2, APOL4, TRIM9, MLLT11, GSE1, CNTNAP2, GNG2, SPAG9, PPFIA2, NRN1, TRIM2, RNF165, NRXN1, CLIP3, C20orf112, RUFY3, DLGAP4, TMEM196, NREP, SV2B, ATP9A, INSM1, DNAJC6, NEUROD1, FAM65A, IRGQ, PTPRR, KIAA1598, DARC, ANK3, KIF21B, INA, UCHL1, ASIC1, HMP19, TUBB2B, ST18, PTCHD2, CNTN2, SEMA6A, SOX11, ERBB4, MFAP2, APC, RUNDC3B, NSG1, MAP2, MAPT, KIF1B, TUBB2A, SPTAN1, NHLH2, CKB, GALNT18, NEDD4L, PHF20L1, CNR1, ROBO2, KIF3C, LOC100131257, AFAP1, DPP6, PARM1, MYT1, SLC17A6, NFIB, KLC1, RELN, GRIK2, KCNQ1OT1, ANKS1B, CNTNAP5, SEPT3, MYT1L, GNAO1, CNTN1, DYNC1H1, SPTBN1, SH3GL2, SERINC1, MALAT1, NFIA, ORC4, CXADR, DPYSL5, KLF7, ERC2, OPTN, KIDINS220, LOC643406, CLSTN2, VAT1, ASTN2, GPM6A, ABCC9, SHISA9, UGDH-AS1, LOC646214, METTL21A, TMEM212, ANKRD20A9P, ODF2L, L2HGDH, MTSS1, LPP, CHGB, ARHGEF26-AS1, ZIC1, MAB21L3, ZBTB20, REXO1L1, GLIPR1L2, IGFBPL1, LPAL2, UGT8, KCNA1, UNC5D, ABLIM1, DYNC1I1, RALYL, FNBP1, MAB21L1, NEK5, LUC7L3, EXPH5, ZBTB18, FGF5, ORAI2, MBOAT1, SETBP1, TMEM130, TRPM3, GNRHR2, CFLAR, DPYSL4, CDH18, NTRK3, ATP6V0A1, TERF2IP, DLGAP1, LINC00461, CHRM3, LHX1, APP, SRGAP3, KCNH7, XKR9, GRIA2, ARRDC3-AS1* |

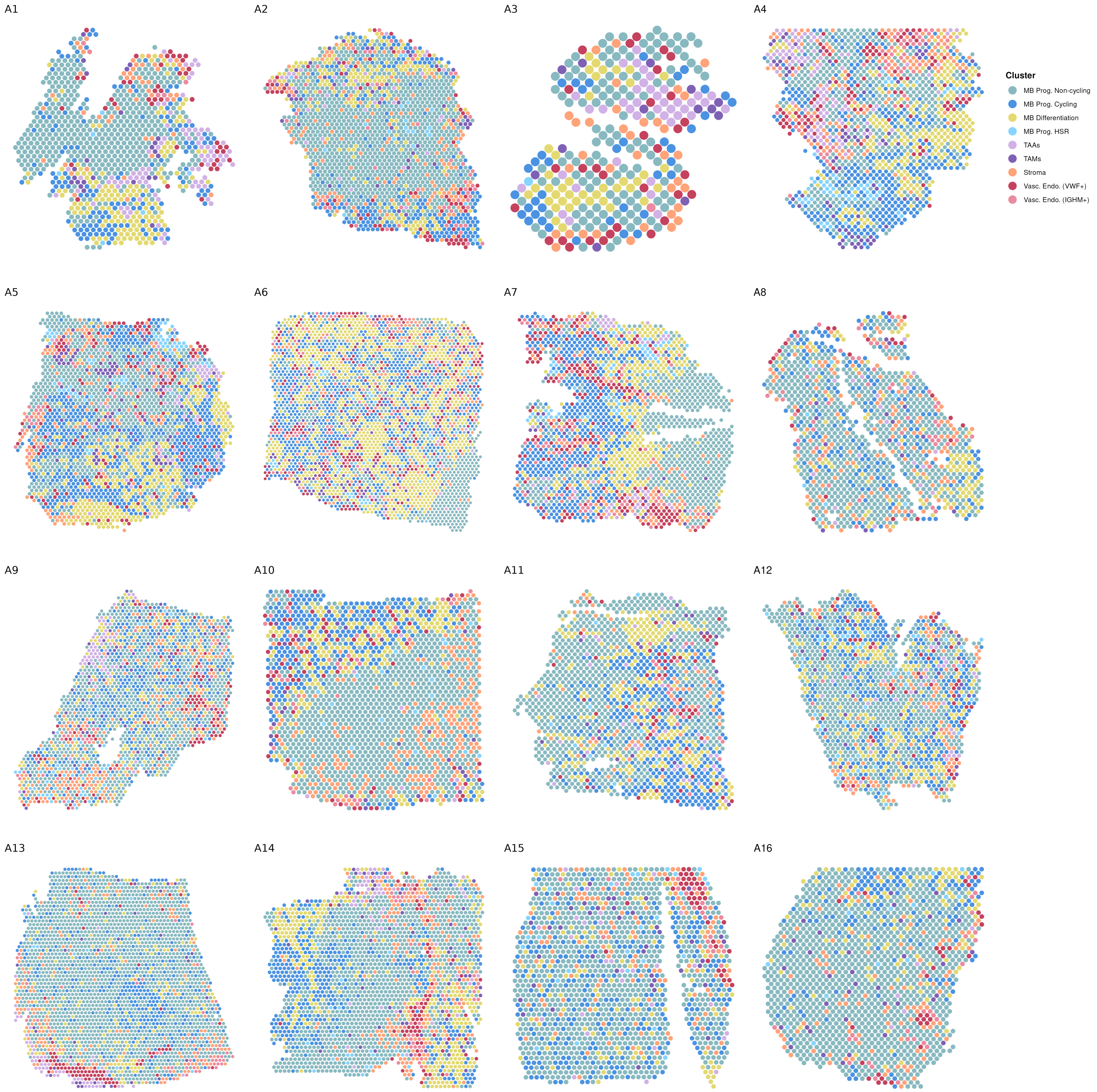


**Figure S1.** Spatial plots illustrating the spatial distribution of clusters across individual samples.

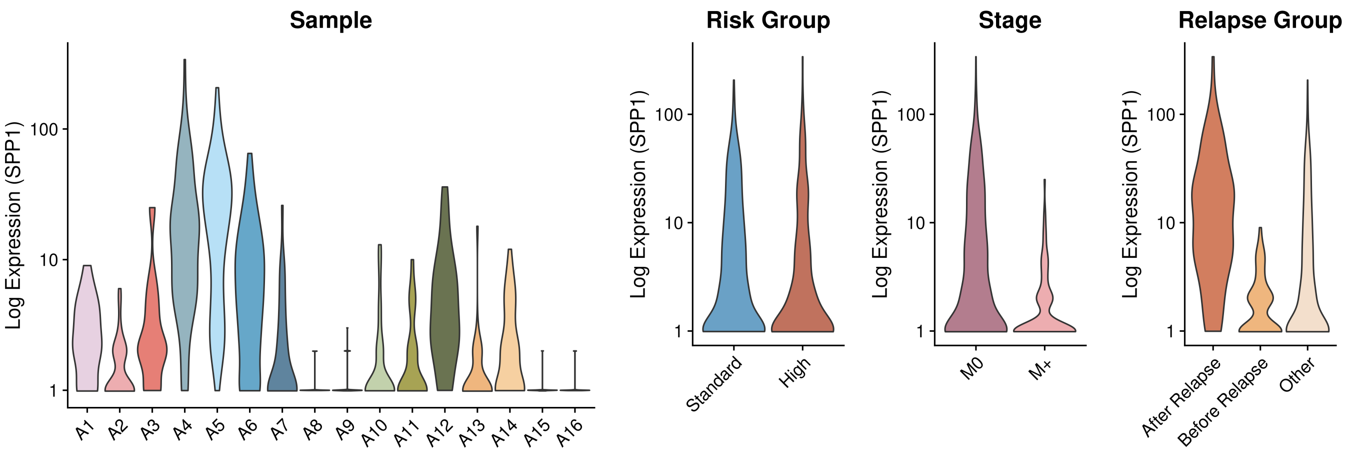


**Figure S2.** Violin plots displaying expression of *SPP1* in voxels from the TAM cluster by sample, risk stratification, metastatic stage, and relapse status.

### **Supplemental Material 2** *Spatial Heterogeneity of the Medulloblastoma Tumor Microenvironment*

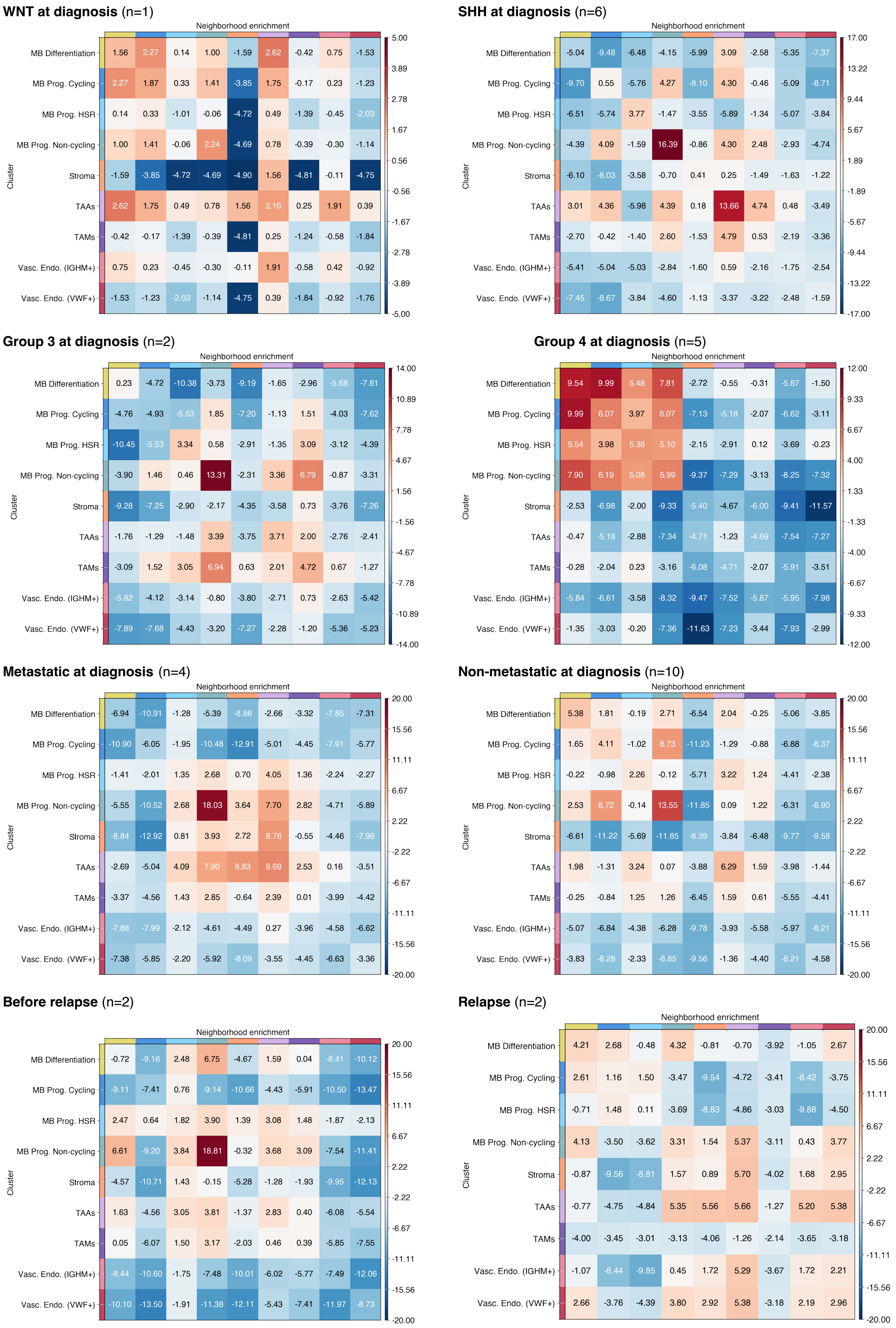


**Figure S3.** Neighborhood enrichment analysis z-score matrices by sample and clinical feature.

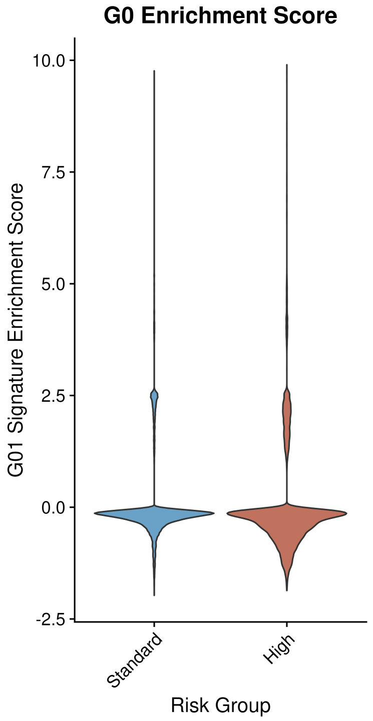


**Figure S4.** Enrichment of G0 signature in HR versus SR samples.
